# IL-10 inhibits STAT1-dependent macrophage accumulation during microbiota-induced colitis

**DOI:** 10.1101/2022.09.01.505766

**Authors:** Izabel Patik, Naresh S. Redhu, Alal Eran, Bin Bao, Anubhab Nandy, Ying Tang, Shorouk El Sayed, Zeli Shen, Jonathan Glickman, James G. Fox, Scott B. Snapper, Bruce H. Horwitz

**Affiliations:** Division of Gastroenterology, Hepatology and Nutrition, Boston Children’s Hospital, Boston, United States; Morphic Therapeutic, Waltham, MA, USA; Computational Health Informatics Program, Boston Children’s Hospital, Boston, United States; Faculty of Veterinary Medicine, Department of Microbiology, Zagazig University, Zagazig, Ash Sharkia, Egypt; Division of Comparative Medicine, Massachusetts Institute of Technology, Cambridge, United States; Department of Pathology, Beth Israel Deaconess Medical Center, Boston, MA, USA; Division of Emergency Medicine, Boston Children’s Hospital, Boston, United States

## Abstract

Loss of IL-10R function leads to severe early onset colitis and in murine models is associated with the accumulation of immature inflammatory colonic macrophages. We have shown that IL-10R-deficient colonic macrophages exhibit increased STAT1-dependent gene expression, suggesting that IL-10R-mediated inhibition of STAT1 signaling in newly recruited colonic macrophages might interfere with the development of an inflammatory phenotype. Indeed *Stat1^-/-^* mice exhibit defects in colonic macrophage accumulation following *Helicobacter hepaticus* infection and IL-10R blockade, and this was phenocopied in mice lacking IFNGR, an inducer of STAT1 activation. Radiation chimeras demonstrated that reduced accumulation of STAT1-deficient macrophages was based on a cell-intrinsic defect. Unexpectedly, mixed radiation chimeras generated with both WT and IL-10R-deficient bone marrow indicated that rather than directly interfering with STAT1 function, IL-10R prevents the generation of a cell extrinsic signal that promotes the accumulation of immature macrophages. These results define essential mechanisms controlling inflammatory macrophage accumulation in inflammatory bowel diseases.

**Summary:** Intrinsic STAT1-function drives the accumulation of macrophages within the colon following the loss of IL-10R signaling. IL-10R prevents this STAT1-dependent process through a non-cell autonomous mechanism.

## Introduction

Rare homozygous loss-of-function mutations in *IL10* or its receptor are among the most common monogenic disorders associated with severe forms of infantile colitis (Glocker et al., 2009, 2010, 2011). Affected patients often fail to respond to conventional therapies, and while bone marrow transplantation maybe curative, there is still a significant unmet need for new and bridging therapies. Despite significant study, a detailed mechanistic understanding of how IL-10 signaling interferes with the development of intestinal inflammation remains elusive. Increasing evidence implicates macrophages as central regulators of intestinal inflammation (Na et al., 2019; Bain and Mowat, 2014). Gut macrophages reside in the lamina propria (LP) beneath the epithelial monolayer and help to maintain mucosal homeostasis under steady state condition. Macrophages within the intestinal LP are phenotypically diverse. At birth, tissue resident macrophages in the colon are primarily embryonically derived (Bain and Mowat, 2014). These macrophages are gradually replaced and continuously replenished by the recruitment of Ly6C^high^ MHCII^low^ blood-derived monocytes, which differentiate into mature Ly6C^-^ and MHCII^high^ macrophages with a tissue resident-like phenotype. Under homeostasis, the recruitment of these Ly6C^high^ monocytes is largely driven by the microbiota and depends on the expression of CCR2 (El Sayed et al., 2022; Caruso et al., 2020; Kang et al., 2019; Redhu et al., 2017; Zigmond et al., 2012).

Mice lacking either IL-10 itself, or either of the two chains of the IL-10 receptor (IL-10R) develop microbiota-dependent colitis, which is associated with a marked increase in the accumulation of inflammatory macrophages in the colon (Redhu et al., 2017; Shouval et al., 2014; Sellon et al., 1998; Kuhn et al., 1993). Remarkably, we and others have shown that mice specifically lacking the IL-10R in myeloid cells (LysM-Cre) also develop colitis characterized by the accumulation of immature Ly6C^+^ macrophages within the colonic LP (Redhu et al., 2017; Zigmond et al., 2012), indicating that IL-10R signaling within the myeloid compartment inhibits the accumulation of immature macrophages. While both newly recruited Ly6C^+^ immature macrophages and Ly6C^-^ mature macrophages are capable of producing pro-inflammatory cytokines following stimuli, it has been argued that newly recruited monocytes are the most significant source of the observable pro-inflammatory cytokine signature including IL-12, IL-23 and ISGs (Interferon-stimulated genes) (Bain et al., 2017; Zigmond et al., 2012), suggesting that the ability of IL-10 to prevent the accumulation of these cells and/or their ability to produce inflammatory cytokines could be an important function involved in inhibiting the development of colitis.

We have recently shown that the loss of CCR2 in mice lacking IL-10R interferes with the accumulation of immature Ly6C^+^ colonic macrophages, likely secondary to the loss of CCR2-mediated monocyte recruitment (El Sayed et al., 2022). However, the identification of intrinsic cytokine response pathways that are necessary for monocyte/macrophage accumulation within the colon of IL-10R-deficient mice is incompletely understood. We have demonstrated that intestinal macrophages isolated from *Il10ra^-/-^* or *Il10rb^-/-^* mice exhibit increased expression of IFN-γ-inducible genes including *Cxcl9, Cxcl11* and *ligp1* (El Sayed et al., 2022; Redhu et al., 2017). Increased IFN-γ production is a hallmark of disease in IL-10R-deficient mouse models, likely as a result of failure to control IL-12 mediated Th1 responses (Ryzhakov et al., 2018; Kullberg et al., 1998). IFN-γ is also identified as a macrophage-activating cytokine, that turns off gene expression associated with M2 type macrophage polarization (Kang et al., 2017; Hu and Ivashkiv, 2009). In addition, patients with defects in IL-10R signaling present with strong IFN-γ expression profiles in colonic tissue (Begue et al., 2011).

STAT1 is the primary transducer of IFN-γ signals (Rauch et al., 2013; Ramana et al., 2000), however the role of STAT1 in driving the development of colitis in IL-10R-deficient mice has yet to be defined. Deletion of STAT1 interferes with the development of DSS-induced colitis and is associated with decreased accumulation of inflammatory macrophages within the colonic LP (Nakanishi et al., 2018), a phenotype also observed in mice lacking *Ifngr1* but not *Ifnar1*. STAT1 has previously been implicated in the development of human IBD (Wu et al., 2007; Mudter et al., 2005; Schreiber et al., 2002), and robust induction of a STAT1-regulated gene network was identified within the terminal ileum of pediatric patients with newly diagnosed Crohn’s disease (CD) (Haberman et al., 2015). It has previously been suggested that IL-10 may directly inhibit STAT1 function and interfere with STAT1-induced gene expression (Ito et al., 1999), but to our knowledge direct evaluation of STAT1 function in regulating monocyte/macrophage accumulation within the colon of IL-10R-deficient mice has not been previously determined. Given the potential promise of JAK/STAT inhibitors in treating inflammatory conditions including IBD, a detailed understanding of STAT1 function in a physiologic model of microbiota-driven colitis could lead to important insights with the potential to guide therapeutic approaches (Cordes et al., 2020; Rogler, 2020; Salas et al., 2020).

Here we show that a cell-autonomous function of STAT1 is necessary for the accumulation of immature macrophages within the intestine of mice lacking IL-10R signaling and is phenocopied by the absence of IFNγR, suggesting that an intrinsic IFNγR/STAT1 pathway drives the accumulation of immature macrophages in mice with defects in IL-10R signaling. Interestingly, IL-10R signaling does not appear to directly interfere with STAT1 function in macrophages, but rather prevents the generation of environmental signals necessary for their accumulation of immature macrophages within the inflammatory microenvironment. These mechanistic studies shed further light on the function of key pathways targeted for therapeutic intervention in IBD.

## Results

### Macrophage accumulation within the colon following blockade of IL-10R signaling requires STAT1

To determine whether STAT1 is necessary for the accumulation of inflammatory macrophages within the colon of mice following blockade of IL-10R signaling, we gavaged parallel groups of age matched WT and *Stat1^-/-^* mice with *Helicobacter hepaticus* (*Hh*) and simultaneously treated them with anti-IL-10RA antibody (Ab) to induce colitis (**Fig. 1A**). Mice were euthanized 16-18 days later. As expected, *Hh*-infected WT mice treated with anti-IL-10RA Ab exhibited histopathological signs of colitis including crypt hyperplasia, abscesses, mononuclear inflammation, and epithelial injury, which resulted in significantly higher histologic activity index (HAI) than in untreated mice (**Fig. 1B and Fig. 1C**). Infection with *Hh* and treatment with IL-10RA Ab in WT mice was also accompanied by increased expression of pro-inflammatory genes (*Il1b, Cxcl2, Il12b*), STAT1-inducible genes (*Cxcl9 and Cxcl11*), and the antimicrobial peptide *Reg3g* within total RNA extracted from combined samples of the cecum and colon (**Fig. 1D**). *Hh*-infected WT mice treated with anti-IL-10RA Ab exhibited a dramatic increase in the absolute number of colonic macrophages (CD45^+^ Ly6G^-^ CD103^-^ CD64^+^ CD11b^+^) compared to sham treated mice (**Fig. 1E and Fig. S1**). In contrast to WT mice, colitis was significantly less severe within the colons of *Hh*-infected STAT1-deficient mice treated with anti-IL-10RA Ab (**Fig. 1B and 1C**). This was accompanied by significantly lower expression of STAT1-dependent genes as well as *Il12b* and *Il1b* within the colon of *Stat1^-/-^* mice compared to WT mice, although we were unable to detect significant differences in the expression of *Cxcl2* and indeed found higher expression of *Reg3g* (**Fig. 1D**). Notably, there was a significant reduction in absolute number of macrophages within the LP (**Fig. 1E**), and this was accompanied by significant reduction in the frequency of immature Ly6C^+^ P1 and P2 macrophages and increases in the frequency of mature Ly6C^-^ P3/P4 macrophages within the total LP macrophage population of STAT1-deficient mice compared to their frequencies in WT mice (**Figs. 1F and 1G**). These results indicate that STAT1 is necessary for the accumulation of immature macrophages observed within the colon of *Hh*-infected mice following blockade of the IL-10R.

**Figure 1.**
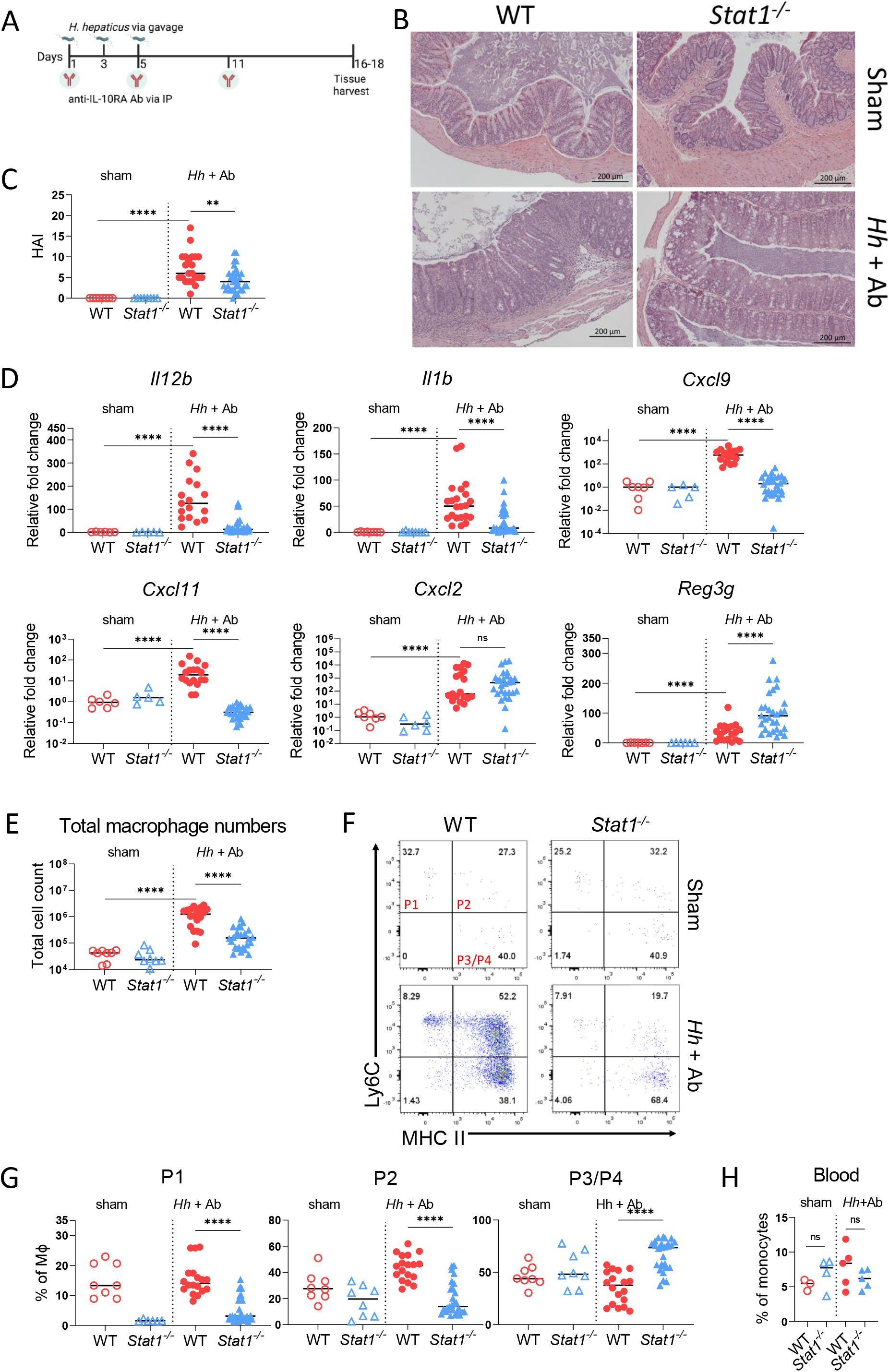
STAT1 is required for colonic macrophage recruitment following infection with *H. hepaticus* and treatment with anti-IL-10RA Ab. **(A)** Schematic showing timing of *H. hepaticus* (*Hh*) infection and anti-IL-10RA Ab administration in model used here. Image created using BioRender. **(B)** Representative histology images (10x) from proximal colonic tissue from sham treated or Hh infected and anti-IL-10RA Ab treated WT and *Stat1^-/-^* mice (***Hh* + Ab**) **(C)** Histopathologic scores of WT and STAT1-deficient mice infected with *Hh* and treated with anti-IL-10RA Ab (*Hh* + Ab), or sham treated with PBS. **(D)** Gene expression by qRT-PCR from colonic tissue. **(E)** Total macrophage numbers from LP of WT or *Stat1^-/-^* mice with or without *Hh* + anti-IL-10RA Ab treatment. **(F)** Representative flow cytometry of MHC II and Ly6C staining of LP macrophages gated on CD45^+^ CD103^-^ Ly6G^-^ CD11b^+^ CD64^+^ cells. **(G)** Percentages of Ly6C^+^ MHC II^-^ (P1), Ly6C^+^ MHC II^+^ (P2) Ly6C^-^ MHC II^+^ (P3/P4) macrophages (MΦ) were determined for mice of indicated genotypes treated with *Hh* and anti-IL-10R Ab or sham treated. **(H)** Percent of circulating monocytes in mice of indicated genotype (CD45^+^/(CD11b^+^ Ly6G^-^ Ly6C^+^ cells)). Results are pooled from minimum three independent experiments, and median values are shown. Statistical significance was determined by nonparametric Mann-Whitney U test. **p<0.01 ****p<0.0001

To determine whether defects in the accumulation of immature macrophages observed in STAT1-deficient mice could be caused by decreased numbers of circulating monocytes, we compared the frequency of circulating CD45^+^ Ly6G^-^ CD11b^+^ Ly6C^high^ monocytes in WT and *Stat1^-/-^* mice either before or following infection with *Hh* and treatment with anti-IL-10RA Ab (**Fig. 1H**). Regardless of inflammatory state of the animals, we were unable to detect significant differences in the frequency of circulating monocytes between WT and STAT1-deficient mice. This result indicates that the reduced accumulation of immature macrophages observed within the colons of *Hh* and anti-IL-10RA treated *Stat1^-/-^* mice is not due to reduced frequency of circulating monocytes.

### STAT1-dependent accumulation of immature macrophages within the colon following blockade of IL-10R signaling depends on IFN-γ

Elevated production of IFN-γ is a hallmark of microbiota-dependent colitis that develops in mice lacking IL-10R signaling, and STAT1 is the primary transducer of IFNγR signaling. This raises the possibility that IFN-γ-induced activation of STAT1 could be required for the accumulation of immature intestinal macrophages observed following blockade of the IL-10R. To address this possibility, we infected *Ifngr1^-/-^* and WT mice with *Hh* followed by anti-IL-10RA Ab treatment as described above (See **Fig. 1A**). Similar to STAT1-deficient mice we found a significant overall decrease in the absolute number of colonic macrophages in IFNγR1-deficient mice compared to WT mice (**Fig. 2A**) as well as decrease in the proportion of the P2 LP macrophage population, and this was accompanied by a reciprocal increase in the proportion of P3/P4 macrophages in IFNγR1-deficient mice (**Figs. 2B and 2C**). Similar to STAT1-deficient mice, the median disease severity score was lower in *Ifngr1^-/-^* mice than in control mice, although this did not reach statistical significance (p=0.12) (**Fig. 2D**). There were significant decreases in the expression of *Il12b, Cxcl9*, and *Cxcl11* within total colonic RNA isolated from IFNγR1-deficient mice compared to WT mice (**Fig. 2E**). Similar to results from STAT1-deficient mice, we did not detect a difference in expression of *Cxcl2* between WT and IFNγR1-deficient mice and expression of *Reg3g* was significantly higher in IFNγR1-deficient mice (**Fig. 2E**). The similar phenotypes observed in mice lacking STAT1 or IFNγR1 suggest that IFN-γ-induced STAT1 signaling is essential for the *Helicobacter*-driven accumulation of immature macrophages in the intestine of mice lacking IL-10R signaling.

**Figure 2.**
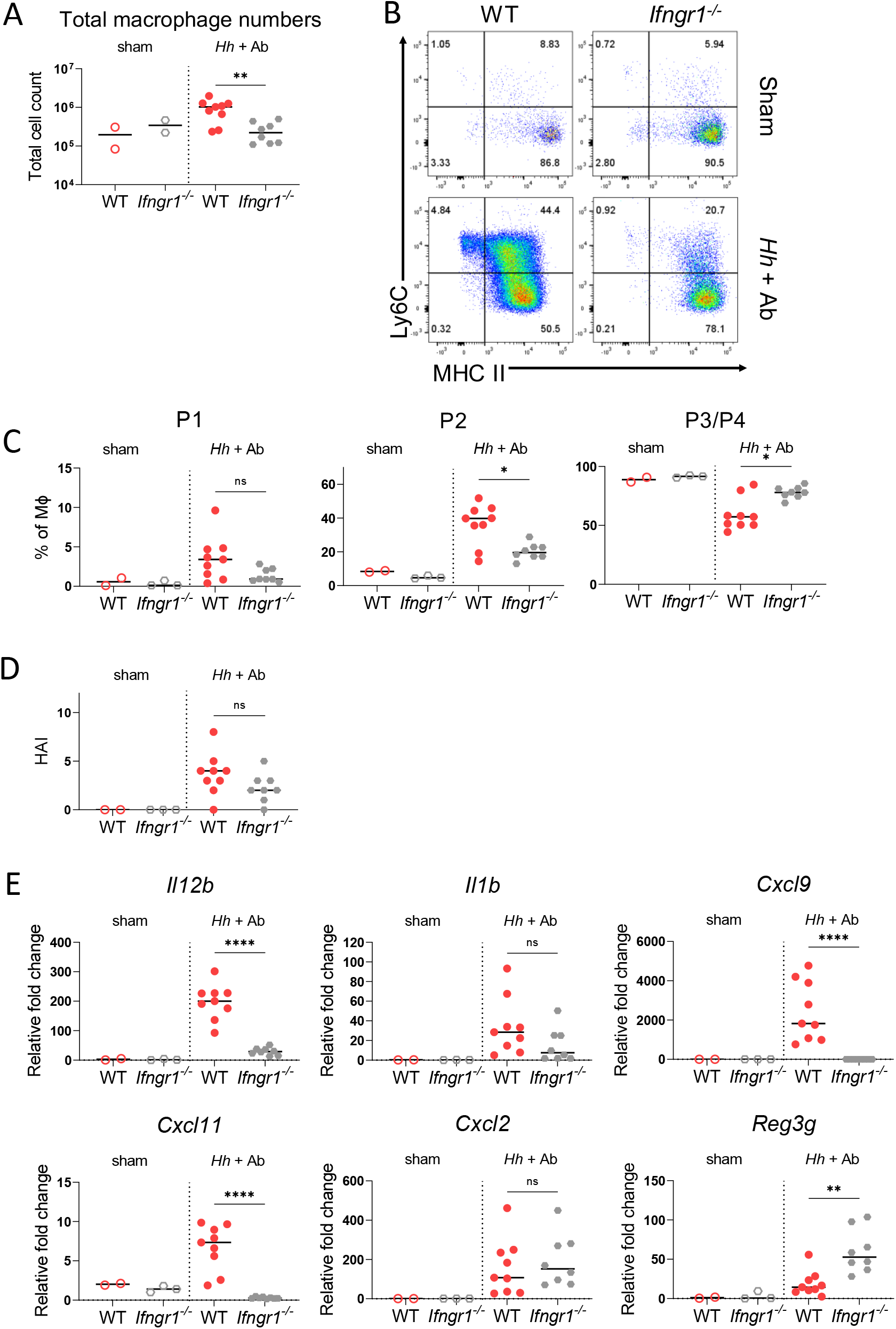
*Ifngr1^-/-^* mice show reduced macrophage accumulation following *Hh* and anti-IL-10RA Ab treatment. **(A)** Total macrophage numbers from the colon of WT or *Ifngr1^-/-^* mice with or without Hh and anti-IL-10RA Ab treatment. **(B)** Representative staining and **(C)** percentages of LP macrophage (MΦ) subpopulations in WT and *Ifngr1^-/-^* mice treated with *Hh* and anti-IL-10RA Ab (*Hh* + Ab) or sham treated with PBS. **(D)** Histopathologic scores of WT and *Ifngr1^-/-^* mice with or without *Hh* and anti-IL-10RA Ab treatment. **(E)** Gene expression by qRT-PCR from colonic tissue. Results are pooled from two independent experiments and median values are shown. Statistical significance was determined by nonparametric Mann-Whitney U test. *p<0.05 **p<0.01 ****p<0.0001

### Loss of STAT1 within the hematopoietic compartment interferes with the accumulation of immature colonic macrophages

To determine whether the function of STAT1 within the hematopoietic compartment is necessary for the accumulation of immature macrophages following blockade of the IL-10R, we generated a series of bone marrow chimeras. In the first set of experiments, we reconstituted lethally irradiated WT (CD45.1) hosts with bone marrow derived from either WT (CD45.1) mice or *Stat1^-/-^* (CD45.2) mice (**Fig. 3A**). Following an eight-week reconstitution period, mice were infected with *Hh* by oral gavage and treated with anti-IL-10RA Ab as above. Mice were euthanized 16-18 days later, and colitis severity and LP macrophages were analyzed as described above. Similarly, to total *Stat1^-/-^* mice, mice that received STAT1-deficient bone marrow exhibited significantly less severe colitis than mice that received WT bone marrow (**Fig. 3B**). The absolute number of macrophages were significantly lower in mice that were reconstituted with STAT1-deficient bone marrow compared to those that were reconstituted with WT bone marrow cells (**Fig. 3C**). Further the frequency of both P1 and P2 macrophages within the donor derived LP macrophage population was significantly lower in mice that were reconstituted with STAT1-deficient bone marrow cells compared to those reconstituted with WT bone marrow cells, while the frequency of P3/P4 macrophages was significantly higher (**Fig. 3D**). Consistent with our previous results, mice that received STAT1-deficient bone marrow cells demonstrated significantly lower expression of *Il12b, Il1b* and *Cxcl9* within total colonic RNA compared to mice that received WT bone marrow following infection with *Hh* and treatment with IL-10RA Ab, although as observed in complete knockouts *Cxcl2* levels remained similar and *Reg3g* expression was significantly higher in mice that received STAT1-deficent bone marrow compared to those that received WT bone marrow (**Fig. 3E**). These results indicated that loss of STAT1 within the hematopoietic compartment interferes with the microbiota-dependent accumulation of inflammatory macrophages following IL-10R blockade.

**Figure 3.**
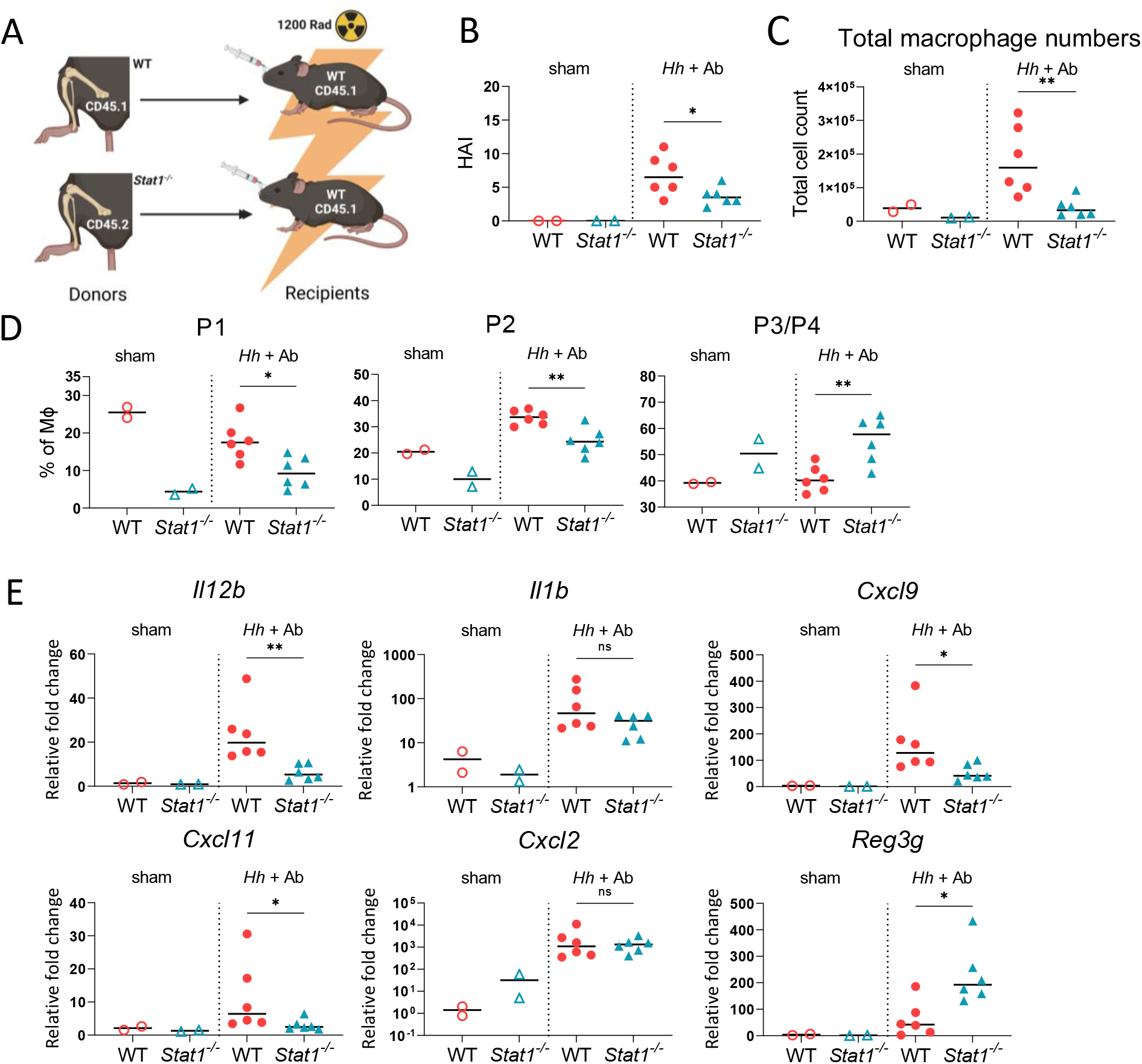
Bone marrow chimeras reconstituted with STAT1-deficient bone marrow show reduced accumulation of macrophages following challenge with *Hh* and anti-IL-10RA Ab. **(A)** Scheme for generation of bone marrow chimeras. WT hosts were reconstituted with either WT or STAT1-deficient bone marrow. Image created using BioRender. **(B)** Histopathologic scores of generated chimeric mice. **(C)** Total macrophage numbers from colon and cecum of chimeric mice with or without *Hh* and anti-IL-10RA Ab treatment. **(D)** Frequencies of colonic macrophage (MΦ) subsets. **(E)** Gene expression by qRT-PCR from pooled colonic tissue. The experiment shows the results from one of two experiments with similar results. Statistical significance was determined by nonparametric Mann-Whitney *U* test. Median values are shown; *p<0.05 **p<0.01.

### Deletion of STAT1 interferes with the development of spontaneous colitis observed in IL10R-deficient mice

We and others have previously shown that loss of IL-10RA in mice carrying the *Cdcs1* colitis susceptibility locus results in the development of spontaneous colitis initiating as early as 3 weeks of age (Redhu et al., 2017). Therefore, to complement the bone marrow chimera experiments described above, we generated an additional set of radiation chimeras in which *Cdcs1* (CD45.1) host mice were reconstituted with bone marrow isolated from either *Cdcs1 Il10ra^+/-^* (CD45.2) (**WT**) mice, *Cdcs1 Il10ra^-/-^ (CD45.2) (**Il10ra^-/-^***) mice, or *Cdcs1 Il10ra^-/-^ Stat1^-/-^* (CD45.2) (**DKO**) mice (**Fig. 4A**). Following reconstitution, mice were housed in conventional SPF conditions for eight weeks to allow the development of colitis and then analyzed. We observed a marked increase in the absolute numbers of donor-derived macrophages within the LP of mice reconstituted with *Il10ra^-/-^* bone marrow cells compared to those reconstituted with control bone marrow cells (**Fig. 4B**). Furthermore, the frequency of P2 macrophages within the donor derived macrophage population isolated from the LP was significantly higher in mice that were reconstituted with *Il10ra^-/-^* bone marrow cells compared to those reconstituted with WT bone marrow cells and we observed a reciprocal decrease in the frequency of P3/P4 macrophages (**Fig. 4C and 4D**). In contrast to mice reconstituted with *Il10ra^-/-^* bone marrow, the median number of donor derived LP macrophages were reduced in mice that were reconstituted with DKO bone marrow cells, and this difference approached statistical significance (p=0.1) (**Fig. 4B**). This was accompanied by a significantly lower frequency of donor derived P2 macrophages and increased frequency of P3/P4 macrophages in the colon of DKO reconstituted mice (**Figs. 4C and 4D**). As expected, histologic scores were quite low in mice that received WT bone marrow and moderately severe in mice reconstituted with *Il10ra^-/-^* bone marrow (**Fig. 4E and Fig. 4F**). We did not observe a significant difference in histologic scores between mice reconstituted with *Il10ra^-/-^* or DKO bone marrow cells (**Fig. 4F**). These results confirm that STAT1 function within the hematopoietic compartment is necessary for the accumulation of immature macrophages observed within the colon of *Il10ra^-/-^* mice.

**Figure 4.**
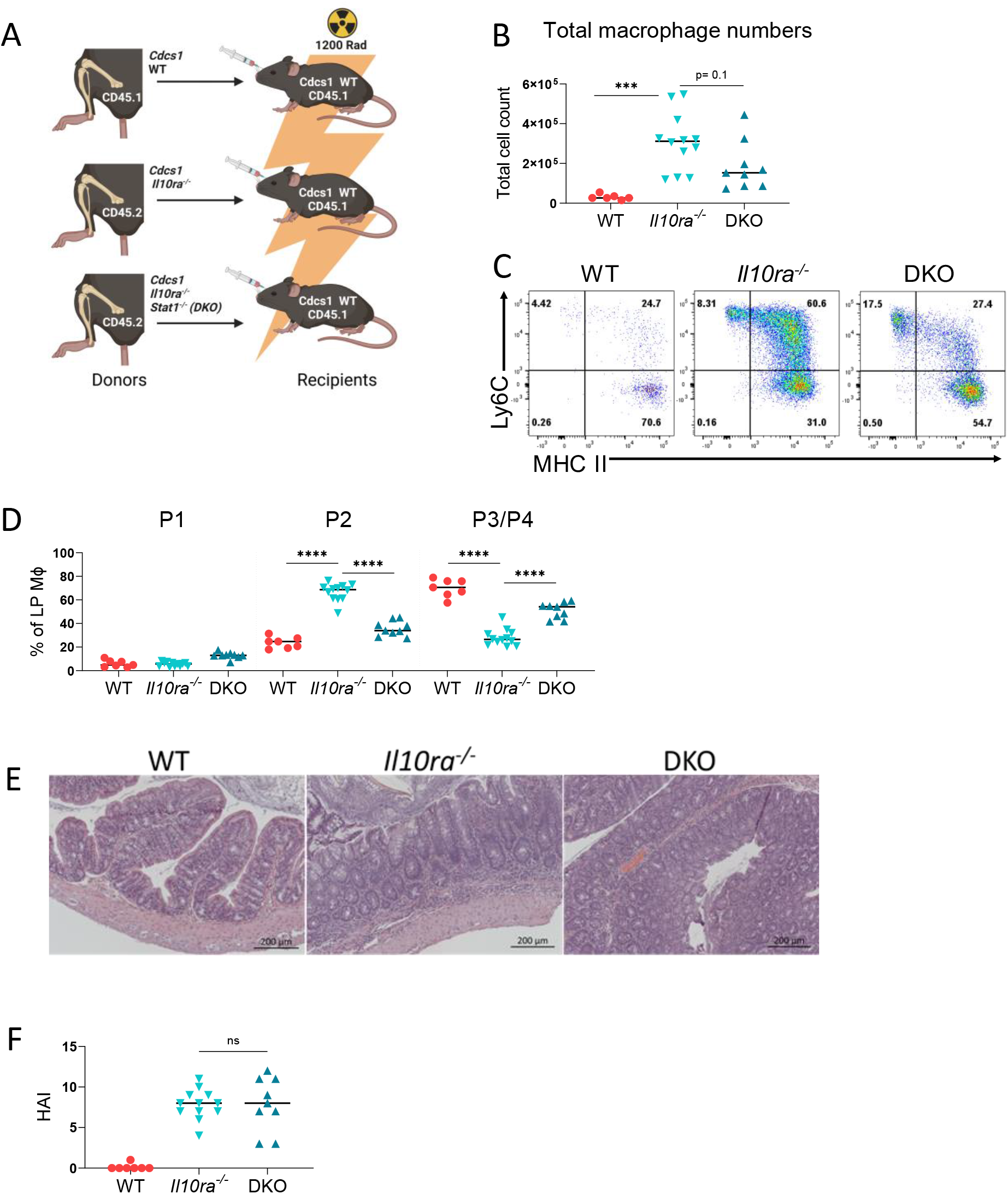
Bone marrow chimeras reconstituted with *Il10ra^-/-^ Stat1^-/-^* bone marrow show reduced spontaneous macrophage accumulation in the LP. **(A)** Scheme of generated bone marrow chimeras. *Cdcs1* (CD45.1) (WT) host were reconstituted with either *Cdcs1* (CD45.1) (**WT**), *Cdcs1 Il10ra^-/-^(CD45.2) (**Il10ra^-/-^***) or *Cdcs1 Il10ra^-/-^ Stat1^-/-^* (CD45.2) (**DKO**) bone marrow cells. Image created using BioRender. **(B)** Total numbers of donor colonic macrophages isolated from chimeric mice. **(C)** Representative flow cytometry showing phenotype of donor-derived colonic macrophages, and **(D)** percentages of macrophage (MΦ) subsets for each group. (E) Representative histology images (10x) from proximal colonic tissue from bone marrow chimeras with H&E staining. **(F)** Histopathological scores for each group. Results are pooled from three independent experiments and median values are shown. ***p<0.001 ****p<0.0001 Mann-Whitney *U* test

### A macrophage-autonomous function of STAT1 is necessary for the accumulation of immature LP macrophages following IL-10RA blockade

As described above, our results indicate that the presence of STAT1 within the hematopoietic compartment is necessary for the accumulation of immature LP macrophages following blockade of IL-10R, suggesting that STAT1 may have a cell-autonomous function that is necessary for macrophage accumulation in the LP. To directly test this hypothesis, we generated mixed radiation chimeras in which irradiated WT (CD45.1) hosts were reconstituted with either WT bone marrow cells, *Stat1^-/-^* (CD45.2) bone marrow cells, or a 1:1 mixture of both (**Fig. 5A and Fig. S2**). Eight weeks later, reconstituted mice were infected with *Hh* and treated with anti-IL-10RA Ab to induce colitis. Sixteen days later, reconstitution was verified by identifying monocytes of the expected genotypes in the blood of these mice. As expected, virtually all blood monocytes in mice reconstituted with WT bone marrow cells were CD45.1^+^ and all monocytes in mice reconstituted with *Stat1^-/-^* bone marrow cells were CD45.2^+^, while in mice reconstituted with a mixture of the two we observed both CD45.1^+^ and CD45.2^+^ monocytes in the blood indicating reconstitution with both WT and *Stat1^-/-^* derived bone marrow cells (**Fig. 5B**). We did note that the proportion in this mixture was skewed towards WT monocytes reflecting either unequal proportions in the initial mixture or perhaps differential reconstitution efficiency.

**Figure 5.**
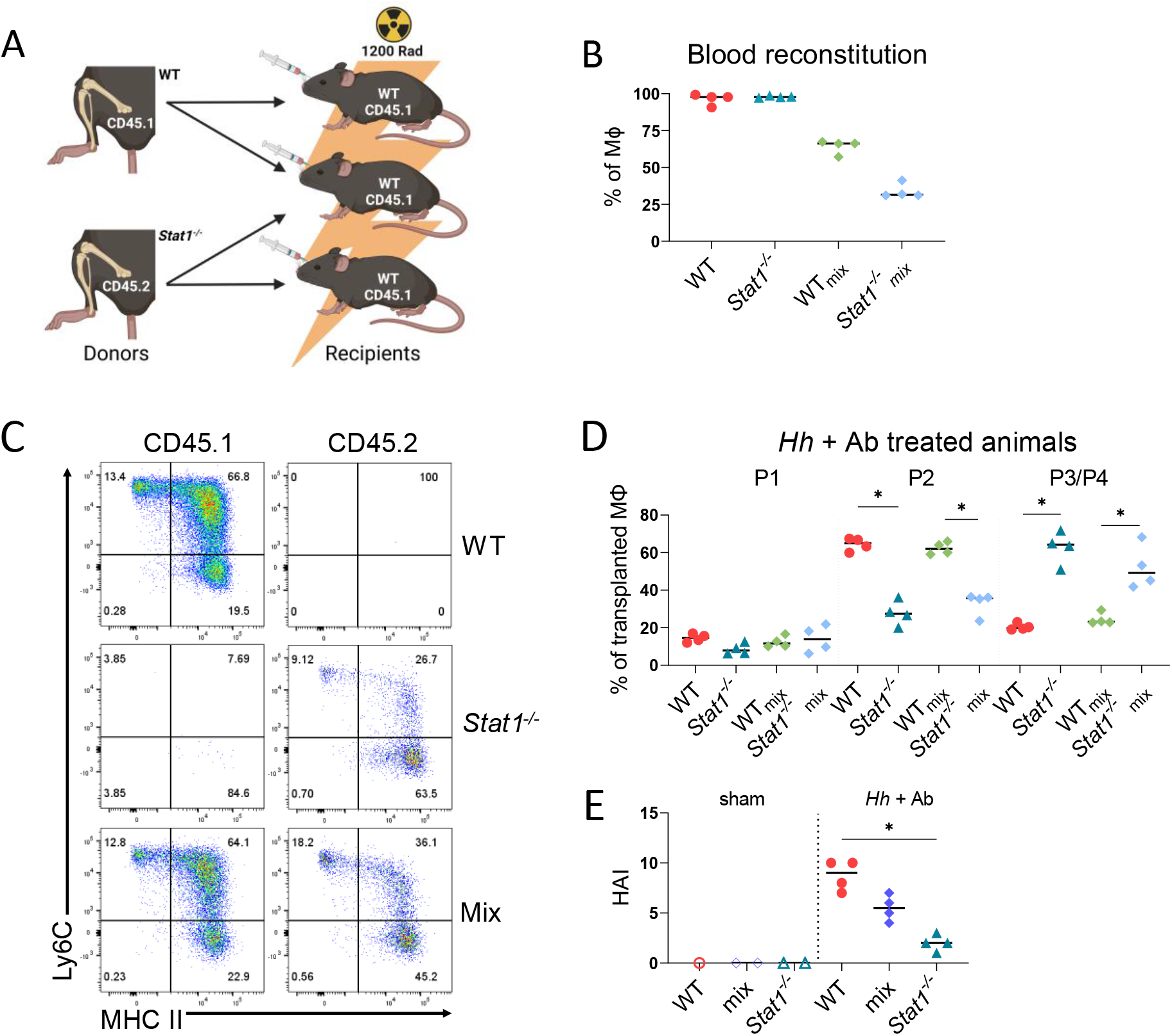
Cell autonomous defect in the accumulation of Stat1^-/-^ immature colonic macrophages following blockade of IL-10R signaling. **(A)** Scheme of generated bone marrow chimeras. WT (CD45.1) hosts were reconstituted with either WT or *Stat1^-/-^* (CD45.2) bone marrow or a 1:1 mixture of both. Image created using BioRender. **(B)** Percentages of CD45.1 and CD45.2 cells that were monocytes in the blood of chimeric mice following induction of colitis. **(C)** Representative flow cytometry of colonic macrophage subsets in chimeric mice. **(D)** Percentages of each macrophage (MΦ) subset within total macrophage populations of each genotype. **(E)** Histopathologic scores from reconstituted mice. Results are from one experiment and median values are shown. *p<0.05 Mann-Whitney *U* test.

We next analyzed the frequency of immature and mature LP macrophages derived from WT or *Stat1^-/-^* bone marrow cells in these chimeras. As expected, based on experiments above, host mice reconstituted with *Stat1^-/-^* bone marrow alone displayed significant reduction in the frequency of donor-derived P2 Ly6C^+^ colonic macrophages compared to hosts that received WT bone marrow cells alone (**Figures 5C and 5D**). Remarkably, in mixed chimeras the frequency of *Stat1^-/-^* P2 macrophages among all *Stat1^-/-^* macrophages in the LP was significantly lower than the frequency of WT P2 macrophages among all WT macrophages, and reciprocally the proportion of P3/P4 macrophages was significantly higher (**Fig. 5D**). Parallel to our previous results from *Stat1^-/-^* mice infected with *Hh* and treated with anti-IL-10RA Ab, mice reconstituted with *Stat1^-/-^* bone marrow developed significantly less severe colitis than mice reconstituted with WT bone marrow (**Fig. 5E**), although the severity of colitis in mice that received mixtures of both was intermediate and not significantly different than the severity in mice that received either WT or STAT1-deficient bone marrow cells alone. These results indicate that the defect in P2 macrophage accumulation observed in the absence of STAT1 is indeed cell autonomous, demonstrating that cell autonomous STAT1 signaling is necessary for the accumulation of immature colonic macrophages observed following blockade of IL-10R signaling.

### IL-10R signaling does not directly inhibit the accumulation of intestinal macrophages, but rather IL-10R-deficient hematopoietic cells confer an inflammatory phenotype on WT macrophages

We have previously demonstrated that specific deletion of IL-10RA in macrophages results in the development of colitis and the accumulation of immature P2 macrophages within the colon (Redhu et al., 2017). Therefore, our observations that STAT1 has a cell autonomous function in macrophages that is necessary for the accumulation of immature macrophages suggests that a critical function of IL-10R signaling could be to directly inhibit STAT1 function.

To directly examine this possibility, we produced a series of radiation chimeras in which *Cdcs1* (CD45.1) (**WT**) mice were lethally irradiated and reconstituted with either WT bone marrow cells, *Cdcs1 Il10ra^-/-^* (CD45.2) (***Il10ra^-/-^***) or a 1:1 mixture of both (**Mix**) (**Fig. 6A**). Eight weeks following bone marrow transplantation mice that received mixtures of WT and *Il10ra^-/-^* bone marrow exhibited the presence of both WT and *Il10ra^-/-^* monocytes within the blood, although the proportion was skewed towards *Il10ra^-/-^*, suggesting as above the possibility of either unequal proportions in the initial mixture or differential reconstitution efficiency (**Fig. 6B**). As anticipated, mice reconstituted with WT bone marrow alone did not develop colitis. Interestingly, mice that received *Il10ra^-/-^* bone marrow cells alone showed clear histologic signs of colitis as did those that received a mixture of WT and *Il10ra^-/-^* bone marrow (**Fig. 6C**). Intestinal LP macrophages in hosts reconstituted with WT bone marrow primarily consisted of P3/P4 macrophages and the frequency of immature P2 macrophage was low, while in hosts reconstituted with *Il10ra^-/-^* bone marrow cells the frequency of donor P2 macrophages was significantly higher (**Fig. 6D and 6E**). Remarkably, the frequency of P2 macrophages derived from either WT or *Il10ra^-/-^* donors mice reconstituted with mixtures of WT and *Il10ra^-/-^* bone marrow cells was virtually identical to the frequency observed in mice reconstituted with *Il10ra^-/-^* bone marrow cells alone, and significantly higher than the frequency of P2 macrophages in hosts reconstituted with WT bone marrow cells alone (**Fig. 6D and 6E**). Because in this system WT macrophages accumulate in the P2 stage even though they have intact IL-10R function, these results imply that IL-10R is unlikely to directly inhibit the accumulation of immature colonic macrophages or the function of STAT1. Rather these data suggest that IL-10R-deficient hematopoietic cells generate a cell extrinsic signal that promotes the accumulation of immature macrophages in the LP through a STAT1-dependent mechanism.

**Figure 6.**
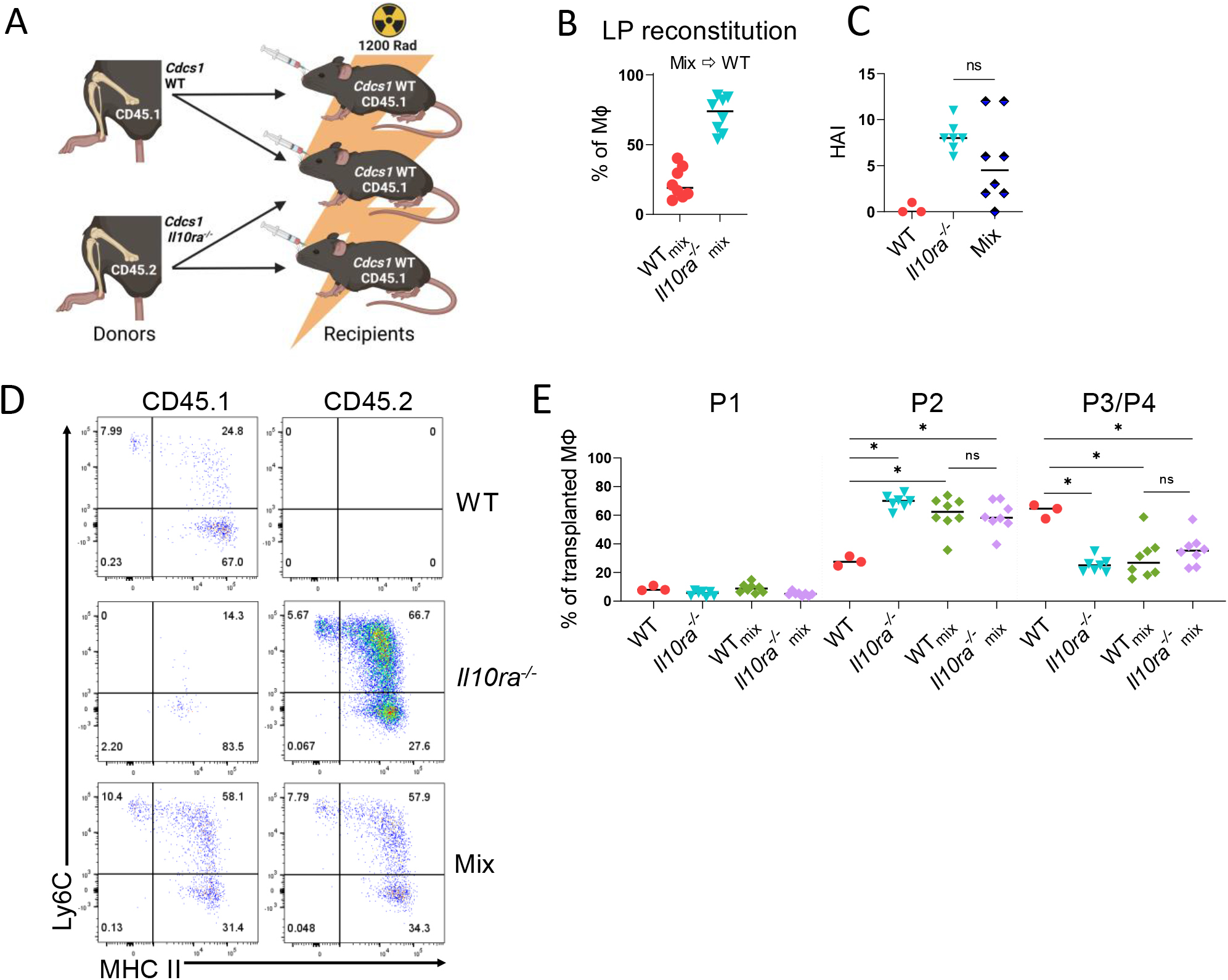
IL-10R deficient hematopoietic cells confer an inflammatory phenotype on WT colonic macrophages. **(A)** Experiment scheme of generated bone marrow chimeras. *Cdcs1* (CD45.1) (**WT**) hosts were reconstituted with either WT or *Cdcs1 Il10ra^-/-^ (**Il10ra^-/-^***) bone marrow, or a 1:1 mixture of both (**Mix**). Image created using BioRender. **(B)** Percentages of macrophages of the indicated genotypes within the blood of mixed chimeras (**WT_mix_** and ***Il10ra^-/-^_mix_*** indicates that cells were isolated from mixed chimeras) **(C)** Histopathologic scores. **(D)** Representative flow cytometry showing macrophage subsets in the colon. **(E)** Percentages of each macrophage (MΦ) subset as a proportion of total macrophages of the indicated genotype. Results are pooled from two independent experiments and median values are shown. *p<0.05 Mann-Whitney *U* test

### Distinguishing cell autonomous and non-autonomous functions of IL-10R signaling in LP macrophages

To further explore the hypothesis that *Il10ra^-/-^* hematopoietic cells influence the phenotype of intestinal macrophages through a non-autonomous mechanism, we isolated LP macrophages from host mice reconstituted with WT bone marrow cells (**WT**), from those reconstituted with *Il10ra^-/-^* bone marrow cells (***Il10ra^-/-^***), and from those reconstituted with 1:1 mixture of both (**WT_mix_** and ***Il10ra*^-/-^_mix_**, respectively), using fluorescence-activated cell sorting. RNA was isolated from these macrophage populations and gene expression analyzed by bulk RNA sequencing. Gene expression profiles were then compared by principal component analysis. WT and *Il10ra^-/-^* LP macrophages clustered independently along both PC1 and PC2. Interestingly, *Il10ra^-/-^* and *Il10ra^-/-^*_mix_ macrophages clustered quite closely together, indicating that the presence of WT macrophages had very little influence on the gene expression characteristics of *Il10ra^-/-^* macrophages. In contrast, WT_mix_ macrophages clustered independently from WT, *Il10ra^-/-^*, or *Il10ra^-/-^*_mix_ macrophages, suggesting that the presence of *Il10ra^-/-^* macrophages strongly influence the gene expression profile of WT macrophages. The differences between WT and WT_mix_ macrophages are most distinguished along PC1, while the differences between WT_mix_ and *Il10ra^-/-^*_mix_, are largely along PC2 (**Fig. 7A**). These results suggest that differences along PC1 are largely driven by cell extrinsic microenvironmental signals generated by the presence of *Il10ra^-/-^* hematopoietic cells, while we suggest that differences along PC2 are determined by genotype (rather than microenvironment) and are therefore cell autonomous.

**Figure 7.**
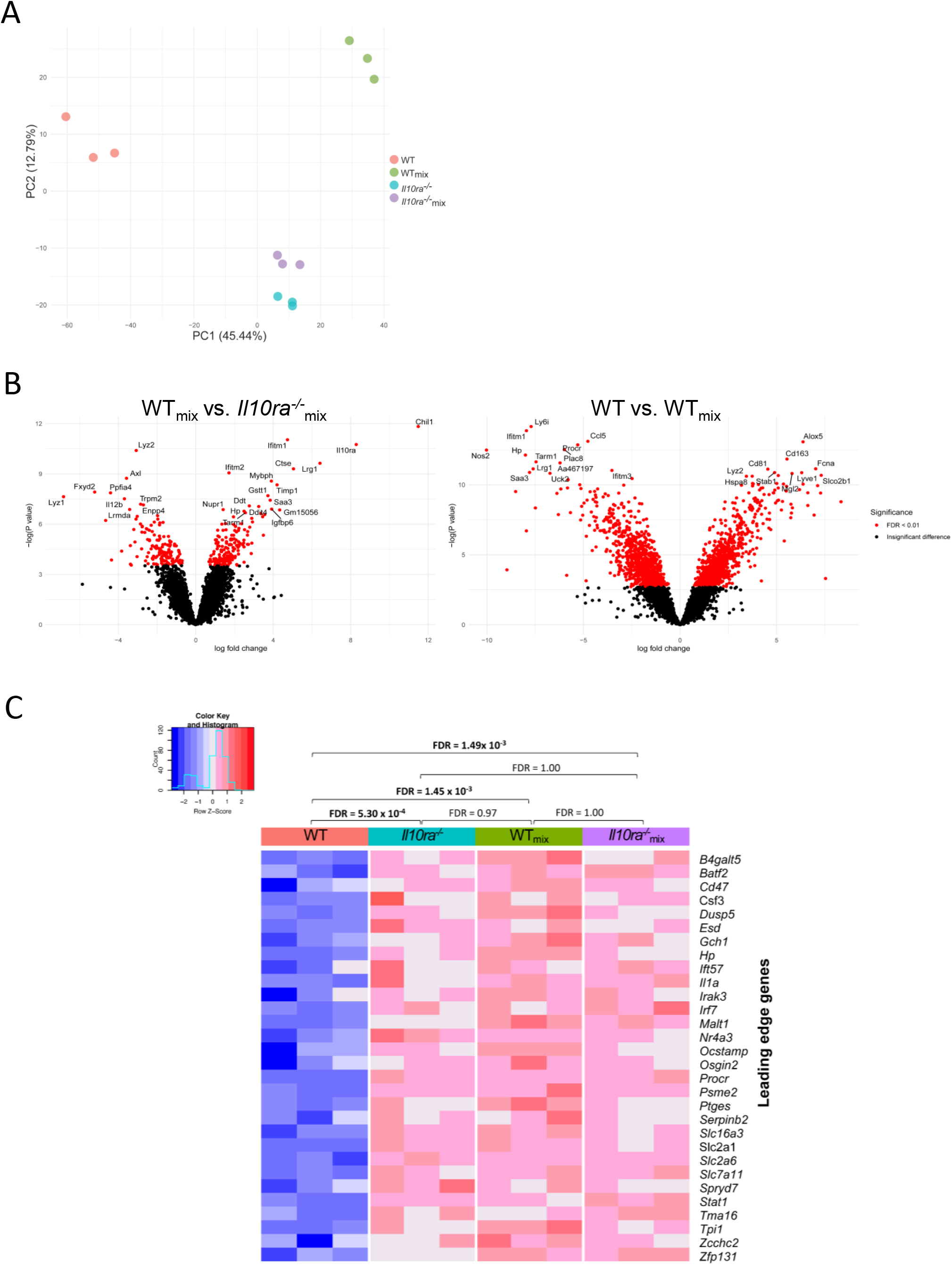
Bulk RNA sequencing from sorted colonic macrophages from mixed bone marrow chimeric mice. **(A)** Principal component analysis of sorted LP macrophages identifies cell autonomous and non-cell autonomous differences in gene expression caused by the absence of the IL-10R. **(B)** Left panel: Comparison of WT_mix_ versus *Il10ra^-/-^*_mix_ identifies genes regulated in a cell autonomous fashion. Genes shown on the left arm of the volcano are expressed at lower levels in WT_mix_ macrophages and those shown on the right arm of the volcano are expressed at higher levels in WT_mix_ macrophages. Right panel: Comparison of WT versus WT_mix_ reveals genes that are regulated in non-cell autonomous fashion by the presence of *Il10ra^-/-^* macrophages. Genes shown on the left arm of the volcano are expressed at lower levels in WT macrophages and those shown on the right arm of the volcano are expressed at higher levels in WT macrophages. **(C)** A heatmap shows the Z-normalized expression of each of 30 leading edge genes from a previously defined gene set of IFN-γ induced genes in bone marrow derived macrophages (Liu et al., 2012) that was found by GSEA analysis to be significantly enriched in comparisons between **WT** and ***Il10ra^-/-^*, WT** and **WT_mix_**, and **WT** and ***Il10ra^-/-^*_mix_** colonic macrophages. Gene set enrichment FDR Q values are shown for the comparison of all experimental groups.

To identify individual genes that were regulated in a cell-autonomous and non-cell autonomous fashion by IL-10RA signaling, we generated a volcano plot of genes differentially expressed between *Il10ra*^-/-^_mix_ and WT_mix_ macrophages, and WT_mix_ and WT macrophages respectively (**Fig. 7B**). We observed that *Il12b* is expressed at lower levels in in WT_mix_ macrophages than in *Il10ra*^-/-^_mix_ macrophages, while the converse is true for *Ddit4*, indicating cell autonomous regulation by IL-10, consistent with our prior observations of rapid inhibition of *Il12b* and induction of *Ddit4* following IL-10 treatment of LPS stimulated bone marrow derived macrophages (Conaway et al., 2017; Ip et al., 2017). In contrast, while it has been suggested that IL-10R signaling might directly induce the development of anti-inflammatory macrophages and suppress the development of inflammatory macrophages, we have observed that *Nos2* (a marker of inflammatory macrophages) expression is lower in WT than in WT_mix_ macrophages, while *Cd163* and *Lyve1* (previously defined markers of mature M2-like colonic macrophages) expression is higher in WT macrophages than in WT_mix_ macrophages (Kang et al., 2019). This suggests that the ability of IL-10R to prevent the development of inflammatory macrophages and promote the development of anti-inflammatory macrophages in the colon is not cell autonomous effect of IL-10R (because IL-10R is present on both of these populations), but rather is the result of differences in the intestinal microenvironment caused by the presence (or absence) of IL-10RA-deficient macrophages.

To further evaluate our hypothesis that IL-10R does not directly regulate IFN-γ-induced STAT1-dependent gene expression in LP macrophages, we performed gene set enrichment analysis (GSEA) focusing on a previously published gene set that is induced in macrophages by treatment with IFN-γ (GSE35825). Our analysis demonstrates that genes in this set are significantly enriched in comparisons between WT and *Il10ra^-/-^* macrophages, WT and WT_mix_ macrophages, and WT and *Il10ra^-/-^*_mix_ macrophages, while no significant enrichment was detected for all other comparisons (*Il10ra^-/-^* vs. WT_mix_, *Il10ra^-/-^* vs. *Il10ra*^-/-^_mix_, or WT_mix_ vs. *Il10ra*^-/-^_mix_) (**Fig. 7C**). This confirms that macrophages with intact IL-10R function (WT_mix_) exhibit elevated IFN-γ/STAT1-dependent gene expression in the presence of *Il10ra^-/-^* macrophages, consistent with our conclusions above that IL-10R signaling does not directly inhibit STAT1-dependent gene expression, but rather inhibits STAT1-dependent gene expression and the accumulation of immature P2 macrophages through a non-cell autonomous mechanism.

## Discussion

Based on prior data from our laboratory and others we had hypothesized that suppression of STAT1 signaling in macrophages was an essential function of IL-10 necessary to inhibit the accumulation of immature macrophages within the LP. Here we have demonstrated that deletion of STAT1 results in marked decrease in the accumulation of immature macrophages within the LP of *Hh*-infected mice following IL-10R blockade, and a significant reduction in the histologic signs of colitis. A similar phenotype was identified in mice lacking IFNγR1. Production of bone marrow radiation chimeras demonstrated that the absence of STAT1 within hematopoietic cells was sufficient to cause a defect in immature macrophage accumulation. Furthermore, using a strategy in which irradiated hosts were reconstituted with mixtures of both WT and STAT1-deficient bone marrow cells, we demonstrate that reduced accumulation of STAT1-deficient macrophages within the LP is the result of a cell-autonomous phenotype. Finally, while these results supported the hypothesis that suppression of STAT1 function by IL-10 is necessary to inhibit immature macrophage accumulation, evaluation of mixed chimeras reconstituted with mixtures of WT and IL-10R-defecient bone marrow cells demonstrated enhanced accumulation of immature WT macrophages that displayed elevated expression of STAT1-dependent genes. These results indicate that IL-10R does not directly suppress immature macrophage accumulation and STAT1-dependent gene expression but rather suppresses these functions in a non-cell autonomous fashion. These observations significantly expand our understanding of the mechanisms through which IL-10 signaling inhibits intestinal inflammation.

Our observation that cell autonomous function of STAT1 is required for the accumulation of immature macrophages within the LP of mice that lack effective IL-10R signaling is consistent with previous results showing reduced accumulation of immature macrophages in STAT1-deficient mice compared to WT mice following treatment with DSS (Nakanishi et al., 2018). The observation that STAT1 is required for macrophage accumulation in mice lacking IL-10R signaling suggests that the dependence of immature macrophage recruitment on STAT1 extends to microbiota-driven models of disease and further is not bypassed even when IL-10R signaling is compromised. This could have important implications for therapy of human disease in situations where macrophage accumulation is a prominent feature. A key unanswered question is how intrinsic STAT1 function supports the accumulation of immature intestinal macrophages. We have not formally investigated the dynamics of STAT1-dependent accumulation of immature macrophage observed in mice following IL-10R blockade, but accumulation could potentially be the result of increased recruitment, *in situ* expansion, decreased cell death, or reduced differentiation into mature macrophage subsets. Although we observed an increase in the proportion of Ly6C^+^ MHCII^+^ LP P2 macrophages compared to other subsets, the absolute number of LP macrophages is markedly elevated following IL-10R blockade, and this extends to all subsets (P1-P3/P4) suggesting that a STAT1-mediated block in differentiation cannot fully explain the macrophage accumulation observed in these animals. A caveat of these studies is that our mouse models did not include the CX3CR1-GFP transgene that is often used to distinguish between maturing P3 and fully mature P4 macrophages (Zigmond et al., 2012), thus we do not know with certainty whether the increase in total macrophages observed in mice following IL-10R blockade extends to fully mature resident-like macrophages, although a prior study using similar methodology did find an increase in the absolute number of P4 LP macrophage following anti-IL-10R Ab treatment of *Hh* infected mice (Bain et al., 2017). Increased percentage of circulating monocytes does also not explain our observations, as we have not observed differences in the percentage of circulating monocytes following IL-10R blockade. Thus, additional studies that include detailed evaluation of LP macrophage dynamics will be required to fully understand the basis for STAT1-dependent macrophage accumulation in the LP of mice lacking IL-10R signaling.

STAT1 is the primary signal transducer of IFN-γ-mediated signaling, and indeed mice lacking IFNγR1 essentially phenocopied STAT1-deficient mice following anti-IL-10R Ab treatment and *Hh* infection. This observation coupled with prior observations regarding the role of IFNγR in macrophage accumulation during DSS colitis strongly suggests that IFN-γ induced STAT1 signaling plays a central role in macrophage accumulation observed in murine colitis (Nakanishi et al., 2018; Kullberg et al., 1998). Nonetheless, we were quite surprised that despite the marked reduction in macrophage accumulation observed in both STAT1- and IFNγR1-deficient mice following IL-10R blockade, we observed levels of residual histologic signs of colitis that remained elevated over untreated control mice. This is not inconsistent with prior studies showing quite variable effects of IFNγR blockade or deletion on the development of colitis in mice lacking IL-10R signaling, although to our knowledge prior studies have not simultaneously evaluated LP macrophage accumulation in parallel with colitis (Kullberg et al., 1998). We suggest that there are several potential explanations for this observation. It is well documented that IFN-γ/STAT1 signaling inhibits expansion of Th17 cells and it is possible that Th17-mediated pathology could compensate for reduced macrophage induced inflammation observed in the absence of STAT1 (Bernshtein et al., 2019; Cypowyj et al., 2012; Liu et al., 2011). Indeed, macrophage-specific deletion of IL-10RA in RAG-deficient mice resulted in enhanced IL-17 responses (Li et al., 2015) and we have also observed increased Th17 responses both in *RAG1^-/-^Il10rb^-/-^* mice following adoptive transfer of unfractionated T cells (Shouval et al., 2014) as well as in humans with IL-10R-deficiency (Shouval et al., 2017). Interestingly, disease in IL-10R-deficient RAG mice following T cell adoptive transfer is abrogated when T cells lack the IL-1R (Shouval et al., 2016; Li et al., 2015), and IL-10R signaling interferes with both transcription of pro-IL-1b and inflammasome dependent production of mature IL-1b in mouse and human macrophages (Shouval et al., 2016; Li et al., 2015). These results raise the possibility that failure to suppress IL-1b production in macrophages may in part be responsible for enhanced IL-17 responses observed in mice and humans with defects in IL-10 receptor signaling. Our data also shows elevated expression of the IL-22-induced antimicrobial peptide *Reg3g* in the colon of STAT1-deficient mice suggesting increased activation of the Th17 pathway. However, prior experiments in WT mice demonstrate that depletion of IL-17 treated with anti-IL-10R Ab and infected with *Hh* actually exacerbated disease severity (Morrison et al., 2015). Moreover, *Hh* colonization and colitis was amplified in *Il17a^-/-^* mice compared to WT mice (Zhu et al., 2022). Both of which are inconsistent with a compensatory effect of IL-17 in STAT1-deficient mice but does not rule-out compensation from another pro-inflammatory pathway.

Another possibility to explain residual colitis observed in STAT1-deficient mice following IL-10R blockade is that the accumulation of immature macrophages is not required for the development of histological signs of colitis in this model. This is consistent with a recent publication from our group demonstrating that despite marked reductions in LP macrophage accumulation in mice lacking both CCR2 and IL-10R compared to mice lacking IL-10R alone, mice lacking both CCR2 and IL-10R still exhibited significantly elevated histologic colitis scores compared to WT mice, indicating that interfering with the accumulation of immature macrophages in the intestine only has a limited role in regulating the histologic manifestations of colitis (El Sayed et al., 2022). This potentially limited role for intestinal macrophages in regulating the severity of colitis seems at odds with strong evidence that macrophage-specific loss of IL-10R signaling is sufficient to drive the development of disease. One potential possibility to explain this phenomenon is that while loss of IL-10R signaling on resident LP macrophages is sufficient to activate inflammatory pathways that drive disease, the accumulation of newly recruited immature macrophages into the colon observed in the absence of IL-10R signaling is not absolutely required for disease development.

Regardless of their exact role in driving histologic signs of disease, the observation of marked STAT1-dependent accumulation of immature macrophages in the absence IL-10R signaling suggested that direct inhibition of STAT1 function by IL-10R could be necessary to prevent immature macrophage accumulation. It has previously been suggested that STAT3, the primary transducer of IL-10R signaling can directly interfere with STAT1-induced gene expression in macrophages by sequestering STAT1 and preventing formation of DNA binding STAT1 homodimers (Ho and Ivashkiv, 2006), although in contrast we have previously argued that IL-10 does not have direct inhibitory effects on IFN-β-induced transcription (Conaway et al., 2017). In fact, data presented here strongly suggest that the ability of IL-10R to inhibit both macrophage accumulation in the colon and the expression of STAT1-dependent genes in colonic macrophages is not cell autonomous, as WT macrophages accumulate and express STAT1-dependent genes at similar levels as IL-10R-deficient macrophages in mixed chimeras between WT and IL-10R-deficient bone marrow cells. This observation strongly suggests that suppression of macrophage accumulation and inhibition of STAT1-dependent gene expression is not a cell autonomous function of IL-10R, but rather more likely an indirect effect caused by suppression of inflammatory mediators within the intestinal environment with the ability to secondarily induce STAT1 activation. The ability of IL-10 to suppress IFN-γ expression within the intestine may be one example of secondary suppression of STAT1 activators. However, the exact pathways through which IL-10 suppresses these secondary mediators remains to be completely defined.

The recognition that IL-10R has potent cell autonomous and non-cell autonomous functions can provide important clues regarding pathways that are directly and indirectly inhibited by IL-10R signaling, and we would propose that understanding these different functions is essential to developing effective therapeutics for children with IL-10R-deficiency and potentially IBD in general. As an example, we and others have proposed that IL-10R signaling may play an important role in generating anti-inflammatory resident macrophages within the intestine (Redhu et al., 2017; Schridde et al., 2017; Girard-Madoux et al., 2016; Shouval et al., 2014). However, our observations that in irradiated host mice reconstituted with mixtures of WT and IL-10R-deficient bone marrow cells, expression of key markers of resident macrophages including *Cd163* and *Lyve1* are markedly reduced in WT_mix_ intestinal macrophages compared to WT macrophages isolated from mice reconstituted with WT bone marrow cells alone, strongly argues that induction of anti-inflammatory resident macrophages is not due to an autonomous inhibitory effect of IL-10R signaling but rather the ability of IL-10R signaling to suppress secondary mediators that interfere with the development of anti-inflammatory resident macrophages in the intestine. In contrast, we have identified a set of genes that do appear to be regulated in a cell autonomous fashion by IL-10R signaling, and these include known targets of IL-10R-mediated suppression including *Il12b*, as well genes induced by IL-10 including *Ddit4*, a key suppressor of inflammasome activation, which we have previously suggested is important to prevent excessive intestinal inflammation (Ip et al., 2017). Based on these observations, we hypothesize that genes regulated in a cell autonomous fashion by IL-10R within intestinal macrophages are essential to prevent the development of colitis, and that further study of this gene set could yield essential clues regarding disease pathogenesis. We suggest that approaches including single-cell RNA-sequencing in mice and comparison of findings using this technique between mice and patients lacking the IL-10R are essential to unravel the key cellular aspects of IL-10R signaling necessary to prevent the development of microbiota-driven colonic inflammation.

## Materials and methods

### Mouse strains

All transgenic and wild-type mice presented here have a C57BL/6 background. C57BL/6J, *Stat1^-/-^* (B6.129S(Cg)-*Stat1^tm1Dlv^*/J), *Ifngr1*^-/-^ (B6.129S7-*Ifngr1^tm1Agt^*/J) and CD45.1 (B6.SJL-Ptprc^a^ Pepc^b^/BoyJ) mice were purchased from Jackson Laboratories and housed under specific pathogen-and viral antibody-free conditions. All other strains used in this study were maintained under specific pathogen-free conditions. *Cdcs1 Il10ra^-/-^* was generated as we have previously described (Redhu et al., 2017). *Cdcs1* (CD45.1) mice were generated by crossing *Cdcs1* (B6.C3Bir-*Cdcs1* (BC-R)) (Bleich et al., 2010) and CD45.1 mice. *Cdcs1 Il10ra^-/-^Stat1^-/-^* mice were generated by crossing *Cdcs1 Il10ra^-/-^* and *Stat1^-/-^* (B6.129S(Cg)-*Stat1^tm1Dlv^*/J) mice. Both male and female mice were used throughout the experiments, and wherever it was possible we used littermate controls. At least two weeks before the initiation of an experiment bedding was mixed to standardize microbiota. Protocols for breeding, housing and experiments were approved by the Animal Resources at Children’s Hospital, according to the Institutional Animal Care and Use Committees (IACUC, Assurance number: A3303-01)

### Colitis induction by *Helicobacter hepaticus* and anti-IL10RA antibody

8 weeks old *Helicobacter*-free mice were gavaged with 200 μl (OD 1.5) *H. hepaticus* inoculums (ATCC 51449 (re-isolated from mice with colitis)) on days 1, 3 and 5. In parallel, 0.5 mg anti-IL-10RA mAb (1B1.3A, Bio X Cell) was administered via IP injection in 150 μl PBS on days 1, 5 and 12. Age-matched littermate controls were included, and sham treated with PBS. Mice were euthanized and tissue was collected for further analysis.

### Lamina propria cell isolation

Cecal and colonic LP cell preparation was carried out as previously described (El Sayed et al., 2022; Redhu et al., 2017). Briefly, colon and cecum were harvested and stripped from epithelial cells by agitating tissue in HBSS media (Gibco) containing 10 mM EDTA and 4.3 mg/ml DTT for 30 min at 37°C. This was followed by collagenase VIII (Sigma) digestion for 35-45 min at 37°C. Remaining undigested tissue was homogenized by repeatedly passing through a 10 ml syringe. Cell suspensions were filtered, washed with PBS, and kept on ice for further use.

### Histology and histopathological analysis

Disease severity was assessed using a histology activity index (HAI) score evaluating H&E-stained sections obtained from the cecum, proximal-, mid- and distal colon. Scoring was based on the severity of mononuclear inflammation (0-4), crypt hyperplasia (0-4), epithelial injury (0-4) and neutrophilic inflammation/crypt abscesses (0-4). Each tissue sample was scored separately for all four metrics by one of the authors (J.G). The reported HAI score is the sum of the individual component scores.

### Microscope image acquisition

H&E stained proximal colonic tissue slides were imaged using a Zeiss Axio Imager Z2 upright microscope at a 10x magnification and files were processed by Zen 3.1 (blue edition) (Carl Zeiss Microscopy GmbH)

### Flow cytometry

All cells were run using a BD LSRFortessa™ Cell Analyzer (BD Biosciences) and analyzed using FlowJo v10.8 (Treestar). For blocking non-specific binding, we used TruStain FcX™ (anti-mouse CD16/32) antibody (BioLegend). Representative examples for the gating strategies are provided in *Supplementary figures 1 and 2*. For LP macrophage analysis cells were gated as follows: SSC-A/FSC-A, FSC-A/FSC-H, FSC-A/FSC-W, Live, CD45^+^ or separated to CD45.2^+^ and CD45.1^+^, CD103^-^, Ly6G^-^, CD64^+^, CD11b^+^, Ly6C/MHC II.

### RNA isolation and quantitative RT-PCR

Tissue from cecum, distal, mid-, and rectal colon were collected and preserved in Trizol reagent (Gibco). Total RNA was extracted following the manufacturer’s protocol, and cDNA generation was carried out using 1 μg total RNA and the TaqMan™ Reverse Transcription Reagents (Invitrogen) following the manufacturer’s protocols. Quantitative RT-PCR was performed using SsoAdvanced Universal SYBR Green Supermix (Bio-Rad) or TaqMan™ Universal Master Mix II, with UNG (Applied Biosystems™) on a QuantStudio™ Flex 6 System and custom primers. Primer sequences are available upon request. Expression was normalized to *Gapdh* or *beta actin* expression, and differences between samples were calculated using the 2^-ΔΔ cycle threshold^ method. Fold-change is reported relative to a sham or uninflamed control.

### Bone marrow chimeras

Recipient mice were irradiated (Gamma Cell 40, ^137^Cs) with a split dose of 1200 Rads 4h apart. The following day bone marrow was harvested from the hind leg bones of donor mice by flushing with sterile PBS. Bone marrow cells were counted, and 4 × 10^6^ cells/mouse were injected into irradiated recipients via retro-orbital injections in 150 μl sterile PBS. Drinking water was supplemented with trimethoprim/sulfamethoxazole (Aurobindo Pharmaceuticals) for 3 weeks (240 mg/ 250 ml water). Eight weeks after transfer, blood was collected via retro-orbital sampling under anesthesia, stained with CD45.1 and/or CD45.2, and analyzed by flow cytometry to evaluate for successful reconstitution.

### Bulk RNA sequencing and analysis

5000-25000 P3/P4 LP macrophages from bone marrow chimeras 8 weeks following reconstitution were FACS-sorted (gated on: CD103^-^, Ly6G^-^, CD11b^+^, CD11c^int^, CD64^+^, Ly6C^-^, CD45.1^+^ or CD45.2^+^) directly into RLT buffer (Qiagen) on a FACS Aria II (BD Life Sciences) and mRNA was extracted using the RNeasy Micro kit (Qiagen). Libraries were prepared using Roche Kapa mRNA HyperPrep strand specific sample preparation kits from 200ng of purified total RNA according to the manufacturer’s protocol on a Beckman Coulter Biomek i7. The finished dsDNA libraries were quantified by Qubit fluorometer and Agilent TapeStation 4200. Uniquely dual indexed libraries were pooled in an equimolar ratio and shallowly sequenced on an Illumina MiSeq to further evaluate library quality and pool balance. The final pool was sequenced on an Illumina NovaSeq 6000 targeting 40 million 50bp read pairs per library at the Dana-Farber Cancer Institute Molecular Biology Core Facilities.

Sequenced reads were aligned to the UCSC mm10 reference genome assembly and gene counts were quantified using STAR (v2.7.3a) (Dobin et al., 2013). Gene counts were then normalized using the trimmed mean of M-values (TMM) approach in edgeR (v 3.15) (Robinson and Oshlack, 2010). Differential gene expression testing was performed using limma (v3.15) (Ritchie et al., 2015), blocking on individual mice.

Since all mice in this study were *Cdcs1* knockouts, in differential gene expression analysis we excluded all *Cdcs1* proximal genes, which might have escaped the targeted knockout. Specifically, we ignored all genes found within 1Mb up and downstream of the expanded *Cdcs1* locus, as previously defined (Bleich et al., 2010). Moreover, since WT mice in the differential gene expression studies were male, and mice of all other groups (*Il10ra^-/-^*, WT_mix_, and *Il10ra^-/-^*_mix_) were female, we excluded sex dimorphic genes in all comparisons involving WT macrophages. Specifically, we ignored genes found to be sexually dimorphic in at least three murine cell types as previously reported (Lu and Mar, 2020). Bulk RNA-seq data generated as part of this study are deposited in the National Center for Biotechnology Information Gene Expression Omnibus (GEO) under the accession number GSE211841. Access available upon request.

We further used gene set enrichment analysis (GSEA; v 4.2.3) (Subramanian et al., 2005) to examine concordant dysregulation of immunologic signatures (C7) of the Molecular Signatures Database (MSigDB) v7.5.1 between groups. In this analysis we removed *Il10ra* and its proximal genes (within 1Mb of its 9q locus), to ensure that *Il10ra* per se and potentially unequal crossover around its locus do not bias set and pathway level convergence.

### Reagents and antibodies

Distributors and origin of reagents, equipment and materials used in this study are indicated in the methods section.

Reagents and antibodies used for flow cytometry and cell sorting: Zombie Violet™ fixable dye (BioLegend), CD45 (clone 30-F11, BioLegend), CD45.1 (clone A20, BioLegend), CD45.2 (clone 104, BioLegend), CD103 (clone 2E7, BioLegend), Ly-6G (1A8, BioLegend), CD64 (clone X54-5/7.1, BioLegend), CD11b (clone M1/70 BioLegend), CD11c (clone N418, BioLegend) Ly-6C (clone HK1.4, BioLegend), MHC II (clone M5/114.15.2, BioLegend).

### Statistical analysis

Statistical analyses were performed using GraphPad Prism 9 unless stated otherwise. P< 0.05 was considered statistically significant and asterix were used to indicate significance as follows: *p< 0.05, **p<0.01, ***p<0.001. ****p<0.0001. Multiple testing correction was achieved using the Benjamini-Hochberg procedure, ensuring that the overall false discovery rate of this study is below 0.05 (Benjamini and Hochberg, 1995).

## Supporting information

Patik et al 2022 Supplementary data

## Author contributions

All authors read and approved the manuscript. I. Patik designed and performed the experiments, analyzed the data, and wrote, reviewed, and edited the paper. N. S. Redhu. designed and performed the experiments, analyzed the data, and reviewed and edited the manuscript. A. Eran analyzed RNA sequencing data and reviewed and edited the manuscript. B. Bao, A. Nandy, Y. Tang performed experiments and reviewed the manuscript. S. El Sayed performed experiments, edited, and reviewed the manuscript. Z. Shen performed the experiments and reviewed the manuscript. J. N. Glickman performed the histology scoring and reviewed the manuscript. J.G. Fox and S. B. Snapper evaluated the data and performed a critical review of the manuscript. B. H. Horwitz designed the experiments, analyzed the data, wrote, reviewed, and edited the manuscript.

## Acknowledgements

We are grateful to Jared Barends, Zach Herbert and Sandra M. Frei for technical assistance. We also thank the IDDRC Cellular Imaging Core, funded by NIH P50 HD105351. We would also like to thank the Beth Israel Deaconess Medical Center Histology Core Facility for processing of histology samples.

This research was supported by a Crohn’s and Colitis Foundation Senior Research Award to B. H. Horwitz and by grants from NIHP30-ES002109, P01CA28842 to J. G.Fox. S. B. Snapper is supported by the National Institute of Diabetes and Digestive Kidney Diseases of the National Institutes of Health under [award number P30DK03485 and RC2DK122532], the Wolpow Family Chair in IBD Treatment and Research, the Translational Investigator Service at Boston Children’s Hospital, and the Children’s Rare Disease Cohort (CRDC) Study.

S.B. Snapper declares the following interests: scientific advisory board participation for Pfizer, BMS, Hoffman La Roche, Lilly, IFM Therapeutics, Merck, and Pandion; grant support from Pfizer, Novartis, Takeda, Amgen; consulting for Hoffman La Roche, Merck, Takeda, and Amgen.

The authors have no additional financial interests.

## Abbreviations

HAI: histologic activity index
*Hh*: *Helicobacter hepaticus*
LP: lamina propria
MΦ: macrophages

