## Supplementary material for "IL-10 inhibits STAT1-dependent macrophage accumulation during microbiota-induced colitis": Patik et al 2022 Supplementary data

**Supplementary Figure 1. Gating strategy to identify lamina propria macrophages in *Stat1*$^{-/-}$ or *Ifngr1*$^{-/-}$ mice.** Single cell suspensions from lamina propria cells were first gated on single cells before gating on live cells. Lamina propria macrophages were identified as CD45$^{+}$ CD103$^{-}$ Ly6G$^{-}$ CD11b$^{+}$ CD64$^{+}$ and cell populations were separated based on Ly6C$^{+}$ MHC II$^{low}$ (P1), Ly6C$^{high}$ MHC II$^{high}$ (P2) and Ly6C- MHC II$^{high}$ (P3/P4). Data is showing one representative experiment.

**Supplementary Figure 2. Gating strategy to identify lamina propria macrophages in bone marrow chimera mice.** Single cell suspensions from lamina propria cells were first gated on single cells before gating on live cells. Lamina propria macrophages were identified as CD103$^{-}$ Ly6G$^{-}$ CD11b$^{+}$ CD64$^{+}$. Congenic CD45.1 and CD45.2 staining separated between WT (CD45.1) and different knock out populations (CD45.2). Cell populations were identified based on Ly6C$^{+}$ MHC II$^{low}$ (P1), Ly6C$^{high}$ MHC II$^{high}$ (P2) and Ly6C$^{-}$ MHC II$^{high}$ (P3/P4). Data is showing one representative experiment.

### Supplementary Figure 1

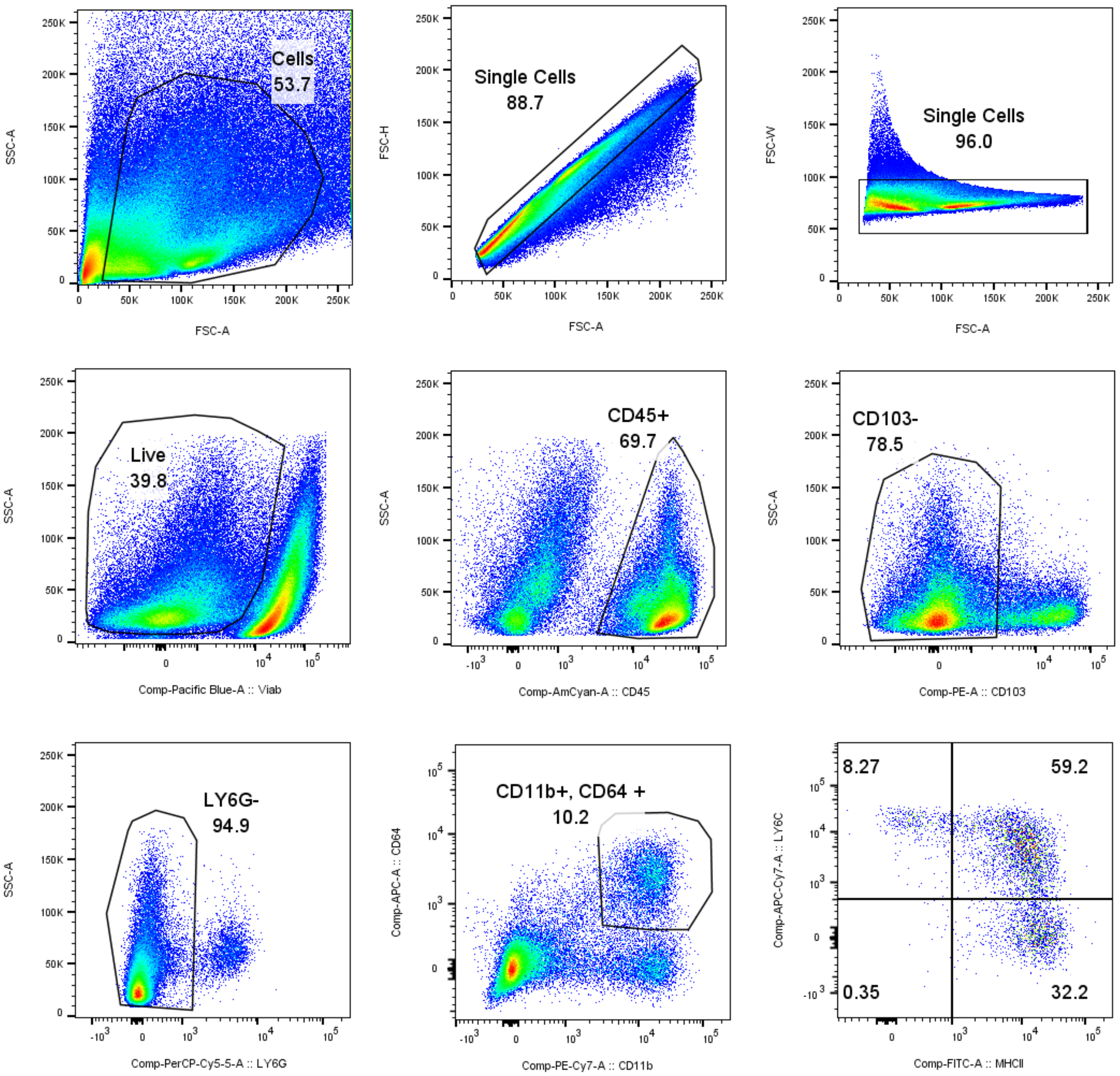

### Supplementary Figure 2

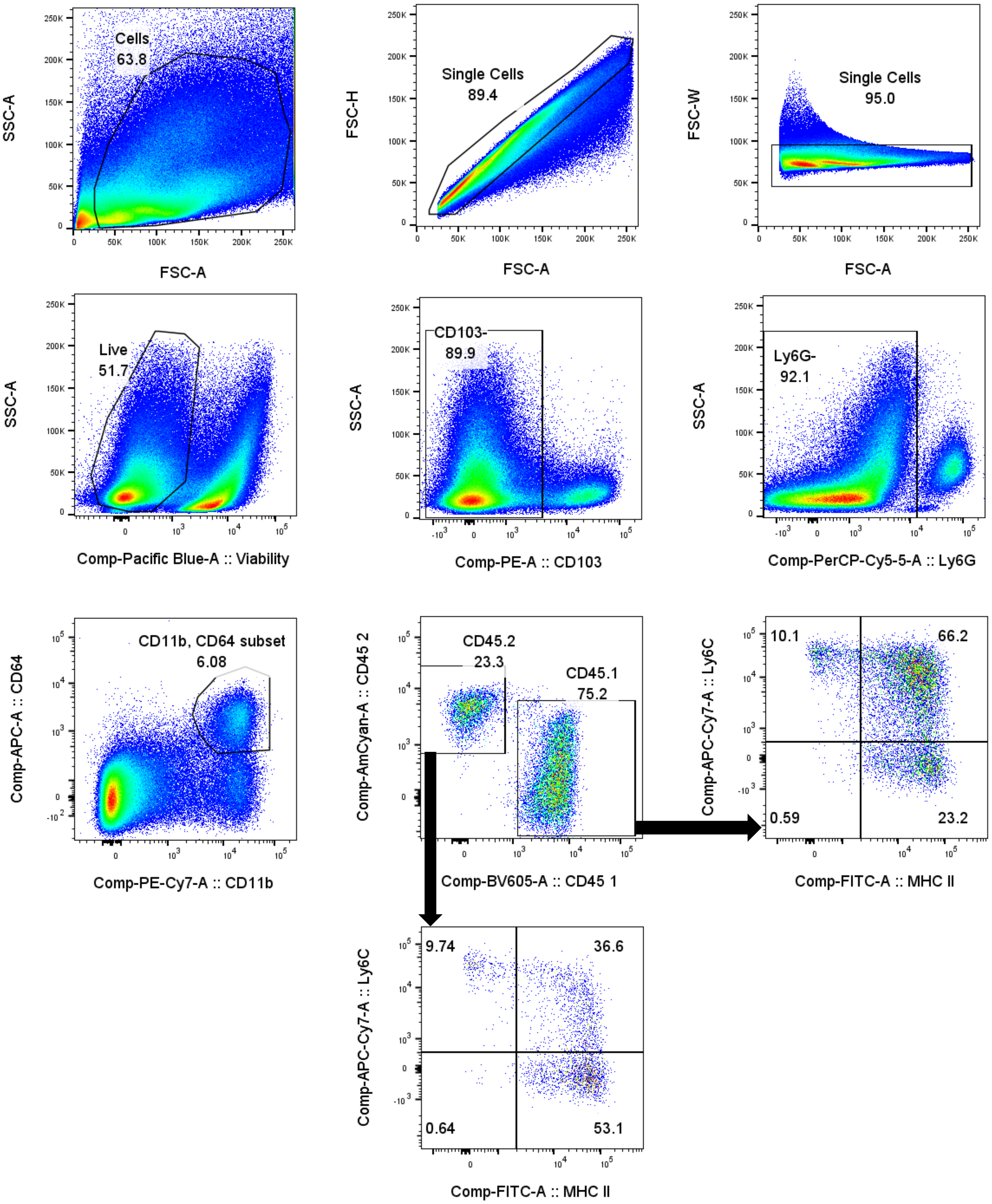
